# Engineered high-density lipoprotein particles that chaperone bioactive lipid mediators to combat endothelial dysfunction and thromboinflammation

**DOI:** 10.1101/2022.02.14.480375

**Authors:** Steven Swendeman, Daniel Lin, Shihui Guo, Alan Culbertson, Andrew Kuo, Michel Levesque, Andreane Cartier, Takahiro Seno, Alec Schmaier, Sylvain Galvani, Asuka Inoue, Samir Parikh, Garret A. FitzGerald, Maofu Liao, Robert Flaumenhaft, Timothy Hla

**Affiliations:** Vascular Biology Program, Boston Children’s Hospital and Department of Surgery, Harvard Medical School, Boston, MA, USA; Division of Hemostasis and Thrombosis, Beth Israel Deaconess Medical Center and Harvard Medical School, Boston, MA, USA; Department of Cell Biology, Harvard Medical School, Boston, MA, USA; Division of Cardiovascular Medicine, Beth Israel Deaconess Medical Center and Harvard Medical School, Boston, MA, USA; Graduate School of Pharmaceutical Sciences, Tohoku University, Sendai, Miyagi 980-8578, Japan; Division of Nephrology, Department of Medicine, University of Texas Southwestern Medical Center, Dallas, Texas, USA; Institute for Translational Medicine and Therapeutics, University of Pennsylvania School of Medicine, Philadelphia, PA, USA

**Author notes:** Current addresses: Sylvain Galvani - Vertex Pharmaceuticals, Boston, MA; Takahiro Seno - Kyoto Prefectural University School of Medicine, Kyoto, Japan.

## Abstract

High-density lipoprotein (HDL) particles suppress inflammation-induced tissue injury via vascular and myeloid cell-dependent mechanisms. As such, HDL-associated bioactive lipids such as sphingosine 1-phosphate (S1P) and prostacyclin (PGI_2_) signal via their respective G protein-coupled receptors on target cells to promote vascular endothelial function and suppress platelet and myeloid-dependent pathophysiology. Here we have constructed a fusion protein of apolipoprotein A1 (ApoA1) and apolipoprotein M (ApoM) (A1M) that forms HDL-like particles and chaperones S1P and Iloprost, stable PGI_2_ analog. The A1M/S1P complex activates S1P receptor-1 (S1PR1) as a Gα_i_-biased agonist and attenuates the inflammation-induced NFκB pathway while A1M/Iloprost acts via IP receptor to inhibit platelet aggregation and promote endothelial barrier function. In addition to enhancing the endothelial barrier, A1M/S1P suppresses neutrophil influx, oxidative burst and inflammatory mediator secretion in a sterile inflammation model. We propose that A1M could be useful as a therapeutic to induce S1P and PGI_2_-dependent anti-inflammatory functions and suppress collateral tissue injury.

## Introduction

Vascular endothelial dysfunction induced by metabolic stress (diabetes, hypercholesterolemia) and autoimmune disease is a major contributor of many chronic diseases (1–3). Restoration of endothelial cell function attenuates inflammation-induced tissue damage and thrombosis and allows efficient resolution process to take place to return to normal homeostasis (4). However, therapeutic strategies to counter endothelial dysfunction remain extremely limited.

Circulating high-density lipoprotein (HDL) particles protect the vascular endothelium. The major structural protein of HDL is apolipoprotein A1 (ApoA1), an amphipathic polypeptide that becomes lipidated after interaction with ABCA1 and G1 transporters (5). Nascent discoidal HDL particles transport cholesterol from tissues to the liver, a process termed reverse cholesterol transport (6). Although reverse cholesterol efflux was thought to be solely responsible for the cardiovascular protective effects of HDL, the ability to attenuate endothelial dysfunction may be equally important (7–10). HDL is associated with numerous proteins and lipids that endow it with cytoprotective and anti-inflammatory properties (11, 12). Among the HDL-associated proteins, the major structural protein ApoA1 decreases cytokine and endotoxin-induced NFκB activation in myeloid cells (macrophages and neutrophils) and the endothelium (13), likely due to suppression of inflammatory receptor signal transduction(14). ApoA1 also enhances endothelial-derived nitric oxide (NO) secretion, which promotes blood flow(15, 16). In addition, HDL suppresses thrombosis by enhancing the activity of prostacyclin (PGI_2_) and attenuation of tissue factor expression(15, 17, 18).

We have characterized apolipoprotein M (ApoM), a member of the lipocalin family of proteins associated with a subpopulation of HDL particles (19). We showed that ApoM chaperones the bioactive lipid, sphingosine 1-phosphate (S1P), a high affinity ligand for G-Protein coupled S1P receptors (S1PRs). The S1P/S1PR1 axis is critical for vascular development, vascular barrier function, NO synthesis, and endothelial survival (20). HDL-bound S1P attenuates cytokine-induced NFκB activation by activating endothelial S1PR1 (21). This signaling axis suppresses lymphopoiesis in the bone marrow (22), endothelial injury in the lung (23) liver fibrosis and promotes hepatocyte regeneration (24). In addition, S1PR1 signaling enables trans-endothelial passage of HDL particles into tissue parenchymal spaces (25). These and other studies suggest that HDL-bound S1P activation of endothelial S1PR1 may be therapeutically tractable to suppress endothelial dysfunction.

We recently reported the development of an ApoM-Fc fusion protein that enhances endothelial barrier function, suppresses ischemia-reperfusion injury of the heart and the brain and immune complex-mediated and acid-induced lung injury (23, 26, 27). However, some of the endothelial protective effects of HDL-S1P are not mimicked by ApoM-Fc/S1P. In this report, we describe the design and characterization of the ApoA1-ApoM fusion protein, which forms spherical nano-sized lipoprotein particles, chaperones multiple bioactive lipids (PGI_2_ and S1P), protects the endothelium, suppresses neutrophil-mediated vascular injury, and platelet aggregation.

## Materials and Methods

### Creation of the ApoA1-ApoM fusion

The ApoA1-ApoM fusion was constructed as follows. Plasmids for murine ApoA1 (Cat# MR203500) and murine ApoM (Cat# MR201811) were obtained from OriGene. The cDNAs for each sequence were amplified to incorporate the following properties:

ApoA1: The endogenous Kozak sequence of ApoA1 and the ApoA1 ORF. The stop codon is replaced with a codon for glycine.

ApoM: A linker sequence coding for one copy of the following five amino acid sequence linker (GGGGS) was added to the 5’-end of ApoM by sequential PCR. The signal peptide of ApoM (AA1-20) was removed and the stop codon was replaced by a glycine followed by a coding sequence to incorporate a 6 x Histidine sequence, to facilitate protein purification, followed by a stop codon.

The two PCR products were linked after NOT1 digestion and the final fusion PCR product was cloned into the pCDH-puro (InvitroGen) expression vector.

Primers used for cloning were:

ApoA1
For: 5’-TTTTCTAGAGAGGAGATCTGCCGCCGCGATCG-3’
REV: 5’-TTTGCGGCCGCGTACGTCTGGGCAGTCAGAG-3’
ApoM
For A: 5’-GGTGGAGGTGGATCT<u>AATCAGTGCCCTGAGCACAGT</u>-3’
Rev A: 5’GATGGTGATGTCCCTTGCTGGACAGCGGGCAGG-3’
For B: 5’-TTTGCGGCCGCTTGGTGGAGGTGGATCT<u>AATCAGTG-3’</u>
Rev B (HIS-TAG) 5’-TTTGGATCCTCAGTGATGGTGATGGTGATGTCCCTTGCTGG-3’

### Expression and purification of the ApoA1-ApoM fusion protein

The resulting pApoA1-ApoM plasmid was transfected into adherent CHO-S cells (InvitroGen) using PEI (Polyethylenimine; 1mM stock solution in water; Sigma). Positive transfectants were obtained by selection in Puromycin (30 μg/ml GIBCO) for 4 days. Cells were tested for expression of the fusion protein by Western blot analysis for ApoA1 (Abcam), ApoM (Abcam) and His-tag (Santa Cruz) expression (data not shown). The drug-selected recombinant CHO-S cells were adapted to serum-free suspension culture using CHO-S medium (CD Forti CHO; ThermoFisher). For large-scale cultures, cells were seeded at 3-5 ×10^5^ cells per ml, and maintained in culture to 2-4×10^6^ cells/ml. Cells were removed from culture by centrifugation at 800xg for 10 minutes at 4°C. The cell culture supernatant was further clarified by ultracentrifugation at >100,000xg for 30 minutes at 4°C. The resulting supernatant was incubated with Ni-Sepharose beads (HisPur Ni-NTA resin, Thermofisher) at a final concentration of 2 ml of beads (packed) /500ml of culture. The slurry was incubated overnight at 4°C and then beads were concentrated by centrifugation at 10,000xg for 5 minutes at 4°C. Beads were washed with 50 volumes of His-wash buffer (20mM Tris ph 8, 400mM NaCl, 10mM Imidizole) until the flow through did not contain detectable protein. Protein was eluted from the column in His-Wash buffer supplemented with 300mM Imidizole. The resulting protein fractions were assayed using Bio-Rad protein reagent and were concentrated with Amicon C15 filters. The final purified protein preparation was analyzed by SDS-PAGE and stained with Coomassie Blue (Bio-Rad; 0.5% suspended in 40% Methanol, 10% Acetic acid.)

### Preparation of S1P, Loading S1P onto ApoA1-ApoM and mass spectrometry analysis

#### Preparation of S1P

1 mg of S1P (Avanti Polar lipids) is resuspended in 13.4 ml of methanol and maintained at 37°C for 12 hours to achieve complete suspension. The solution is dispersed into 134μl aliquots and dried under vacuum for 45 minutes to 1 hour at 37°C. Dried S1P is maintained at −20°C until use.

#### Loading S1P onto ApoM

We had previously described a method for loading S1P onto recombinant ApoM (26). Essentially, purified protein was suspended in PBS at 1 mg/ml (~20μM) and mixed with 160μM S1P by gentle pipetting. The sample is subjected to 3 rounds of 30 second sonication in a bath sonicator and allowed to incubate for >24hours by nutation. The resulting product is subjected to FPLC to separate the free S1P from the ApoA1-ApoM/S1P protein complex and the resulting protein fractions were concentrated using Amicon C15 filters.

### Loading of S1P onto Albumin

S1P was loaded onto Albumin as described (28). Essentially, a 0.4% solution of PBS-Albumin (Fatty acid free, Sigma) was added to dried S1P and subjected to bath sonication for 3x cycles for 1 minute/cycle and maintained at 4°C for 24 hours.

### Negative stain electron microscopy analysis of ApoA1-ApoM

ApoA1-ApoM sample was diluted to 0.02 mg/ml and 2.5 μL was applied to glow-discharged copper EM grids covered with a thin layer of carbon film. Grids were strained with 1.5% (w/v) uranyl formate, blotted, and allowed to air dry. Negatively stained EM grids were imaged on a Tecnai T12 electron microscope (Thermo Fisher Scientific) operated at 120 kV at a nominal magnification of 67,000 x using a 4k x 4k CCD camera (UltraScan 4000, Gatan), corresponding to a pixel size of 1.68 Å. EM images were binned by two (3.36 Å per pixel) and particles were selected and subjected to two-dimensional classification (29).

### Mass Spectroscopy analysis of S1P

Mass spectroscopy-based quantitation of S1P was performed as described previously (30). S1P-bound ApoM or ApoA1-ApoM proteins or plasma from mice injected with these proteins was extracted in Methanol (20:1; Methanol:analyte, (vol/vol).

### Loading of Iloprost onto ApoA1-ApoM chaperone

We employed a standard sonication and thermal cycling method for creating lipidated ApoA1-ApoM as described(31, 32). Either Phosphorylcholine 2mM (Avanti) or a combination of DMPC and DMPG (7:3 molar ratio) (Avanti/Sigma) were resuspended in Chloroform:Methanol (95:5) and distributed into Eppendorf tubes and dried under vacuum. Lipid was resuspended in PBS buffer, heated at 37°C for 5-10 minutes and sonicated (20 seconds) to create a lipid suspension. Purified human ApoA1 (Sigma) or purified ApoA1-ApoM protein was resuspended in PBS (10:1 mol/mol) was added to the lipid slurry and subjected to either continuous bath sonication until the suspension exhibited clarification (Indicating lipidation) or subjected to repeated cycles (3–5) of 37°C heating (5 minutes) followed by 5 minutes of bath Sonication at RT. In each method, Iloprost (Cayman Chemical) was added to the solution at a final concentration of 1mM. The resulting lipidated solution was subjected to FPLC purification (26) and relevant fractions corresponding to recombinant HDL nanoparticles were collected and concentrated on Amicon Filters.

### S1P and Iloprost TEER analysis of HUVEC

Transendothelial Electrical Resistance (TEER) was performed on HUVEC as described previously (26). Essentially, HUVECs were maintained in growth medium (HGM; M199 medium supplemented with 10% FCS (Corning), 1:100 Pen-Strep (Sigma), 8mM Glutamine (Sigma), Heparin (Sigma, 100U, LMW) and endothelial cell growth supplement(33). All cultures were maintained on fibronectin (2 μg/ml) coated plates. For TEER analysis we employed a 96 well ECIS system (96W10idf PET array, Applied BioPhysics). Wells were coated with fibronectin (2 μg/ml) resuspended in saline for 30 minutes at room temp. HUVECs were harvested, resuspended in HGM at a cell density of 25-30×10^3^ cells/well and allowed to adhere overnight. Prior to analysis, culture media was removed and replaced with M199 media supplemented with Pen-Strep/Glutamine and 1% FCS for 30 minutes. For stimulation studies, either Iloprost (Cayman Biochemicals) or Iloprost loaded onto ApoA1 or ApoA1-ApoM chaperones, or S1P prepared on ApoA1-ApoM or ApoM-Fc chaperones were added to cultures and TEER studies were performed from 3-24 hours.

### S1P, Angiopoietin1 (Ang-1) and Activated Protein C (APC) regulation of endothelial barrier function

Angiopoietin1 (R&D systems) was used at a final concentration of 300ng/ml for all studies. Thrombin (Millipore Sigma) was used at a final concentration of 1U/ml for all studies. Activated Protein C was obtained from Enzyme Research laboratories. For initial studies, Ang-1 was evaluated alone and in combination with ApoM-Fc/S1P. For Thrombin studies, Ang-1, ApoM-Fc/S1P, and Thrombin were co-added at the initiation of the study. For APC experiments, APC alone or in combination with A1M/S1P was added to the culture for 1 hour prior to the addition of Thrombin and then TEER analysis was performed for an additional 2-8 hours. Data from TEER studies were statistically analyzed using GraphPad Prism 7 (GraphPad Software, San Diego, CA).

### Nanobit analysis of S1P receptor Activation

The NanoBiT system, which employs split luciferase consisting of small and large fragments (SmBiT and LgBiT, respectively) and was previously utilized to monitor interactions of GPCRs with ß-arrestin or specific Gα/βγ complexes in response to GPCR activation (34) was used to characterize ApoA1-ApoM nanoparticles containing S1P. Briefly, HEK293A cells maintained in DMEM (GIBCO) supplemented with 10% FCS and Pen-Strep, were dispersed into 6 well plates and allowed to adhere overnight. For functional studies, cells were transfected with appropriate combinations of reporter plasmids. S1PR1-SmBiT and LgBiT-beta-arrestin fusion proteins or Gα_i_-SmBiT combined with Gβ_1_ and LgBiT-Gγ_1_ were employed as described previously (34). After 24hrs, cells were harvested, resuspended with the luciferase substrate Coelenterazine (Cayman Chem; 50 μM), dispersed into white opaque-bottom 96-well plates (Greiner) and maintained at RT for 2 hours to quench background. S1P containing samples were added by multi-channel pipetting and the plate was immediately analyzed for luminenescence for 30 minutes in a SpectraMax L 96-well plate reader (Molecular Devices). Data was integrated as described (34).

### Inhibition of TNF-α dependent NF-κB activation

TNFα-dependent NF-κB signaling was determined by using an NFκB-Luciferase based reporter assay system (pGL4.32[*luc2P*/NF-κB-RE/Hygro]; Promega). We created a stable NFκB reporter cell line in human microvascular endothelial cells (HMEC-1; ATCC). HMEC are maintained in 10% FCS/Pen-Strep MCDB media supplemented with 2mM L-Glutamine (Sigma), EGF (2ng/ml; R&D systems) and Hydrocortisone (1ng/ml; Sigma). Using TNFα (10ng/ml; R&D Systems), we performed both dose and temporal analyses to establish optimal induction of the Luciferase response (Data not shown). The 5 ×10^5^ reporter cells were distributed to 12 well plates and allowed to adhere for 24 hours. Media was replaced with MCDB media supplemented with 1% FCS, L-Glutamine. Cells were pre-treated with either media, ApoA1 (Purified Human ApoA1; Sigma), ApoA1-ApoM, ApoA1-ApoM/S1P, ApoM-Fc/S1P, or Albumin-S1P for 10 minutes and 2ng/ml of TNFα. Samples were extracted using cell lysis buffers (Promega), luciferin substrate was added and the plates were measured for luminenescence for 8-15 minutes in a SpectraMax L 96-well plate reader (Molecular Devices).

### Inhibition of TNF-α-induced ICAM-1 expression in HUVEC

HUVECs were plated as described above. After 24 hours cells were shifted to M199 media supplemented with 1% FCS, 8mM Glutamine, 1X Pen-Strep. After 30 minutes, cells were pre-treated for 10 minutes with media, ApoM-Fc/S1P (100nM S1P), Iloprost (100nM), ApoA1-ApoM (200μg/ml), ApoA1-ApoM-Iloprost (200ug/ml; 100nM Iloprost) or ApoA1-ApoM/S1P (200μg/ml100nM S1P) or in combinations and subsequently treated with TNF-α (10ng/ml). After 5 hours, cells were lysed with cell lysis buffer (TBS-T; 20mM Tris-pH8, 160mM NaCl, 1% Triton, 1X protease inhibitor cocktail (Sigma)), collected and centrifuged at 20,000 x g for 5 minutes at 4°C, extracts were analyzed by 10% SDS-PAGE, transferred to nitrocellulose membrane (Bio-Rad), blocked in 5% milk, and probed with antibodies to ICAM-1 (1:1000, sc-8439, Santa Cruz Biotech) and Actin (1:5000, sc-8432, Santa Cruz Biotech). Blots were developed with appropriate secondary antibodies linked to HRP and visualized by chemiluminescence (Immobilon Western, EMD Millipore) using a ChemiDoc Imaging (Biorad).

### Iloprost-induced activation of the CREB–luciferase signaling

In order to evaluate the activity of lipoprotein-bound Iloprost to activate the I prostanoid receptor (IPr), we examined the downstream cAMP-dependent CREB-luciferase reporter system (pGL4.29 [luc2P/CRE/Hygro]; Promega). HEK293T or A cells were maintained in DMEM (InVitrogen) supplemented with 10%FCS and Pen-Strep (Corning). Cells were harvested and dispersed 2-3 x 10^5^ cells/well in 6 well plates and allowed to adhere for 24 hours. Media was replaced and cells were transfected using PEI (Polyethylenimine; 1mM stock solution in water; Sigma) with either 0.3 μg/well of reporter plasmid alone or co-transfected with pCMV-PI, which encoded the IP receptor. After 24 hours, media was replaced and cells were incubated for 8 hours at 37°C with media containing either vehicle, Iloprost (5 nM - 100 nM), or ApoA1-ApoM-Iloprost (1-32μg/ml) by titration. Samples were lysed, luciferin was added and luminenescence was measured for 30 minutes in a SpectraMax L 96-well plate reader (Molecular Devices). Data was integrated as described. Untransfected HEK293T or A cells do not respond to Iloprost stimulation, and gained responsiveness upon IPr expression.

### Inhibition of Human Platelet aggregation *in vitro*

Human studies were approved by the institutional review board of the Beth Israel Deaconness Medical Center (PI: R. Flaumenhaft). Whole blood samples were drawn from healthy donors in the presence of 10% sodium citrate (Sigma, S577-50ml) and were spun at 1000rpm using Beckman centrifuge (GS-6K) for 20min. The top Platelet-rich Plasma (PRP) layer was collected and rested in a 37ºC water bath for 30min. PRP was diluted with one-fifth volume of acid-citrate-dextrose buffer, Prostaglandin E_1_ (PGE_1_) was added to a final concentration of 0.15 μM. The mixture was spun in a conical-bottom tube at 2000 g for 10 min, and the platelet pellet was collected and suspended in pre-warmed HEPES-Tyrode-Glucose Buffer (HTG). The platelet concentration was determined and adjusted to ~2.5×10^5^/μL with HTG on a Hematology cell counter System (Drew Scientific, 850FS).

Light Transmission Aggregometry was used to evaluate the platelet response to agonists and antagonists. In a 4-channel aggregometer (Platelet ionized calcium aggregometer, Chronolog Corp Model 660), human platelets were stirred in a cuvette at 37°C. Human platelets were incubated with A1M/Iloprost, ApoA1/Iloprost or A1M/S1P alone or in combination for ~10min before adding thrombin receptor (PAR-1) agonist SFLLRN peptide (2μM). Data were analyzed using AGGRO/LINK software package Ver 5.2.5 and Microsoft office professional plus 2013.

### Inhibition of Neutrophil Reactive Oxygen Species (ROS) release

A1M, A1M/Iloprosst and A1M/S1P were assayed for inhibition of f-Met-Leu-Phe (f-MLP) dependent Reactive Oxygen Species generation in isolated murine neutrophils as follows. Peritoneal neutrophils were isolated from C57Bl/6 mice after thioglycolate elicitation. Thioglycolate (2ml of 2% v/v) resuspended in water was administered by intraperitoneal injection(35). After 4 hours, mice were euthanized and 5 ml of HBSS was injected intraperitoneally, the abdomen was messaged for 1 minute and the peritoneal fluid was removed. The resulting cells were pelleted, RBCs were lysed with ACK buffer, re-pelleted and resuspended in PBS-glucose. Cells were counted, 5 x10^5^ cells were dispersed into 96 well clear bottom plates containing luminol 100mM and 10 U/ml of horseradish peroxidase and read for blank background. F-Met-Leu-Phe (fMLP; 10 μM) was added and plates were read on a SpectraMaxL1 (Molecular Devices, San Jose, CA) over 5 minutes as described previously. The area under the curve was calculated by using GraphPad Prism 7 (GraphPad Software, San Diego, CA).

### Analysis of Neutrophil Influx in Thioglycolate Induced Sterile Peritonitis

Mice were injected with Thioglycollate as described above. One hour after thioglycolate induction, mice were injected with either PBS (vehicle), or 200μg of either A1M or A1M/S1P. After 4 hours, cells were isolated as described above, lavage fluid was saved for cytokine analysis (see below), and cells were washed, counted, and stained for flow cytometry using the neutrophil marker Ly6G (PE-anti-Mouse Ly6G; Cat No. 127608; BioLegend), gating for side-scatter and PE+ cells.

### Analysis of Cytokine Content of Peritoneal Lavage Fluid in Thioglycolate-Induced Sterile Peritonitis

Approximately 3.5 ml of peritoneal lavage fluid was collected from each mouse after thioglycolate induction (see above). After cell clarification by centrifugation (300g x 5’), 1ml aliquots of fluid was prepared and immediately frozen in Liquid N_2_. 500ul of peritoneal fluid was analyzed for cytokine expression using the Proteome Profiler Mouse Cytokine array Kit, Panel A (Cat No. ARY006; R&D Systems) following the manufacturer’s instructions. After development, blots were subjected to IMAGEJ analysis and data were analyzed using GRaphpad Prism 7 as above.

### Animals

C57Bl/6 mice were obtained from Jackson Labs. All *in vivo* experiments were performed according to approved experimental protocols by IACUC at Boston Children’s Hospital.

## Results

### Design, expression and characterization of ApoA1-ApoM/S1P (A1M/S1P) nanoparticles

Recently, we described ApoM-Fc, a fusion protein of ApoM with the Fc domain of IgG and demonstrated its ability to enhance endothelial cell function *in vitro* and *in vivo* (24, 26, 27). However, unlike purified HDL, ApoM-Fc/S1P did not inhibit cytokine-induced endothelial dysfunction. We therefore hypothesized that a fusion protein of ApoA1 and ApoM might mimic endothelial protective HDL functions better than ApoM-Fc. We designed and constructed the ApoA1-ApoM (A1M) fusion protein with a flexible linker domain (GGGGS) (Figure 1A, Supplementary Figure 1) (36). In addition, we added a 6X-Histidine tag at the carboxyl-terminus to enable rapid protein purification (Supplementary Figure 1). We expressed and purified A1M protein from conditioned media from stably transfected CHO-S cells (Figure 1B).

**Fig. 1.**
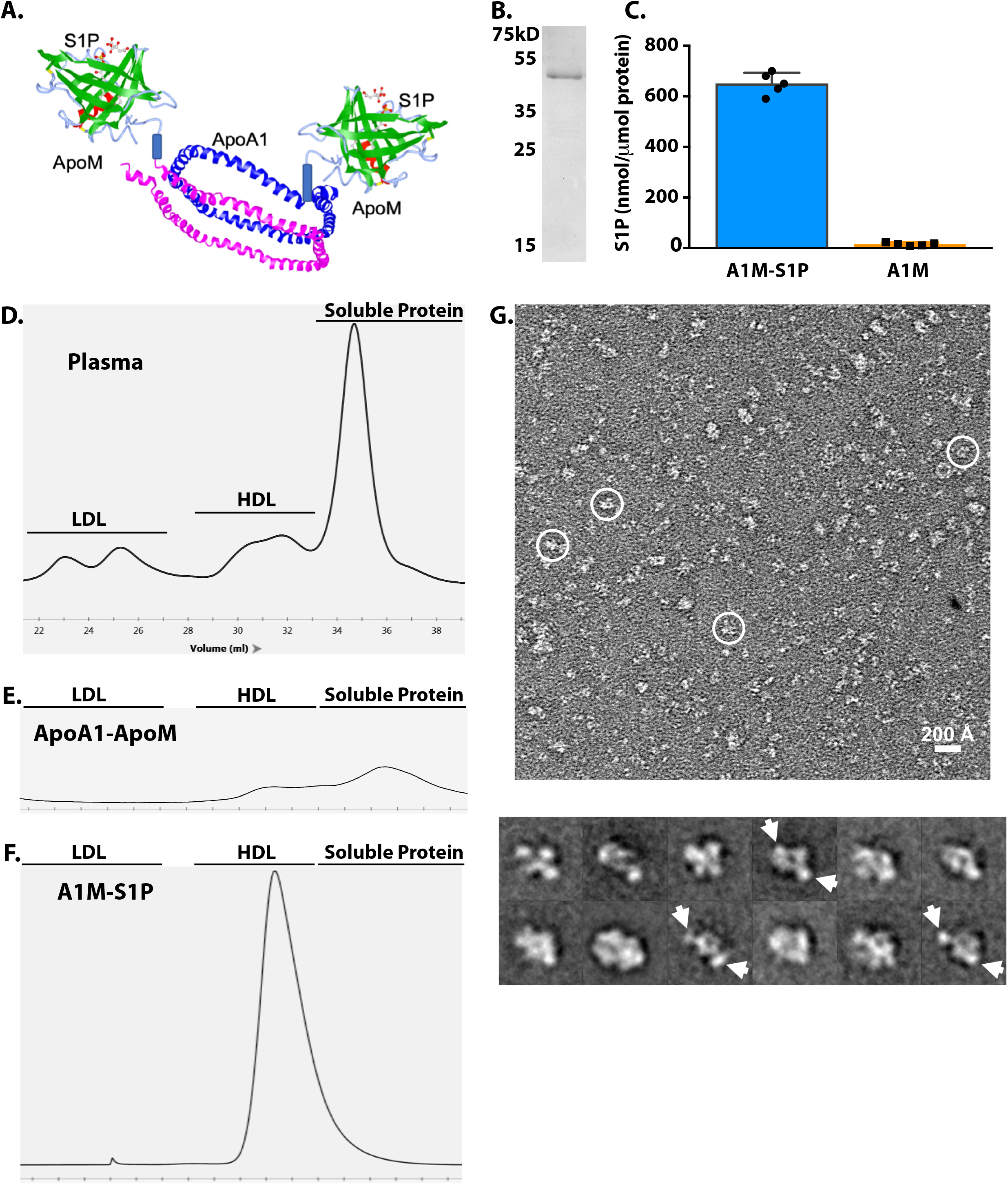
Production, purification, and characterization of S1P binding by the ApoA1-ApoM fusion protein. **(A)** Schematic model of ApoA1-ApoM (A1M). **(B)** Purified A1M (4 μg) was separated by reducing 10% SDS-PAGE and stained with Coomassie brilliant blue. **(C)** Purified A1M (100 μg) was incubated or not with S1P for 24 hours, purified by gel filtration chromatography and analyzed for S1P content by electrospray ionization-MS/MS. The resulting data are the mean + S.D.; N=5. **(D, E, F)** Representative FPLC elution profiles (OD 280) of 200μl of mouse plasma, purified recombinant A1M, and A1M after lipidation and S1P loading. (D) FPLC elution of plasma standards: LDL (22-26ml) HDL (28-32ml), Soluble protein fraction (32-38ml). (E) nascent A1M protein (100μg) (F) lipidated A1M/S1P (1 mg). **G)** Representative negative-stain EM image of A1M with some particles highlighted in white circle (left) and 2D averages with potential ApoM densities indicated by arrows (right). Box dimension of each 2D average is 215 Å.

A1M was loaded with S1P as described previously and purified by FPLC gel filtration chromatography (26). Under these conditions, A1M protein was loaded with ~54-68 mol% of S1P (Figure 1C). In contrast, unloaded purified A1M contained < 0.5 mol% S1P, presumably from endogenous sources (CHO cells or FBS-containing cell culture medium).

Since the ApoA1 moiety of HDL can be lipidated, we sonicated A1M with a mixture of phosphatidylcholine and S1P as described (31). FPLC elution profiles of the samples were compared to a reference of fractionated mouse plasma ((Figure 1D). Analysis of native A1M reveals a mixture of lipidated (~30%) and unlipidated fractions (70%) (Figure 1E). After the lipidation procedure, the A1M loaded with S1P eluted near the HDL region, suggesting efficient conversion to HDL-like particles (Figure 1F). Purified A1M/S1P lipoprotein particles were analyzed by transmission electron microscopy after negative staining. As shown in Figure 1G, many particles are ~ 8-12 nm in diameter with various apparent shapes. This is in line with the conformational heterogeneity of lipoprotein and the flexible linkage between ApoA1 and ApoM.

Several two-dimensional (2D) averages show two side lobes connecting to a central disc-shaped density (Fig. 1G, right), which is consistent with the expected shape of A1M complex (Fig. 1A). These studies suggest that A1M/S1P complex forms nanometer-sized lipoprotein particles that stably bind S1P.

### A1M/S1P induces Gα_i_-biased activation of S1PR1

The ability of the A1M/S1P complex to activate S1P receptors was studied using the real-time NanoBiT system that monitors coupling to heterotrimeric G proteins and ß-arrestin (34). Dose- and time-dependent activation of S1PR1 by A1M/S1P was contrasted with albumin-S1P and ApoM-Fc/S1P. Over a 4-log range of ligand stimulation, similar S1PR1-dependent activation of Gα_i_ (Figure 2A) and β-arrestin coupling (Figure 2B) was observed for all three chaperones complexed with S1P, suggesting that ApoM chaperone function is not impaired by fusion to ApoA1 or nanoparticle formation. However, temporal analysis of receptor stimulation at saturating ligand concentrations (1μM S1P) revealed the Gα_i_-biased nature of ApoM-bound S1P. When bound to S1P, all three chaperones activated Gα_i_ with similar kinetics (Figure 2C). However, β-arrestin activation by chaperone-bound S1P (Figure 2D) occurred later than Gα_i_ activation (T_1/2_ ~ 360 s *vs*. 1,320 s). Moreover, β-arrestin activation of ApoM-bound S1P (A1M/S1P and ApoM-Fc/S1P) was significantly slower than albumin-bound S1P (T_1/2_ ~ 1,320 s *vs.* 960 s; P=0.012). These data are consistent with A1M/S1P and ApoM-Fc-S1P exhibiting Gα_i_-biased activation of S1PR1.

**Fig. 2.**
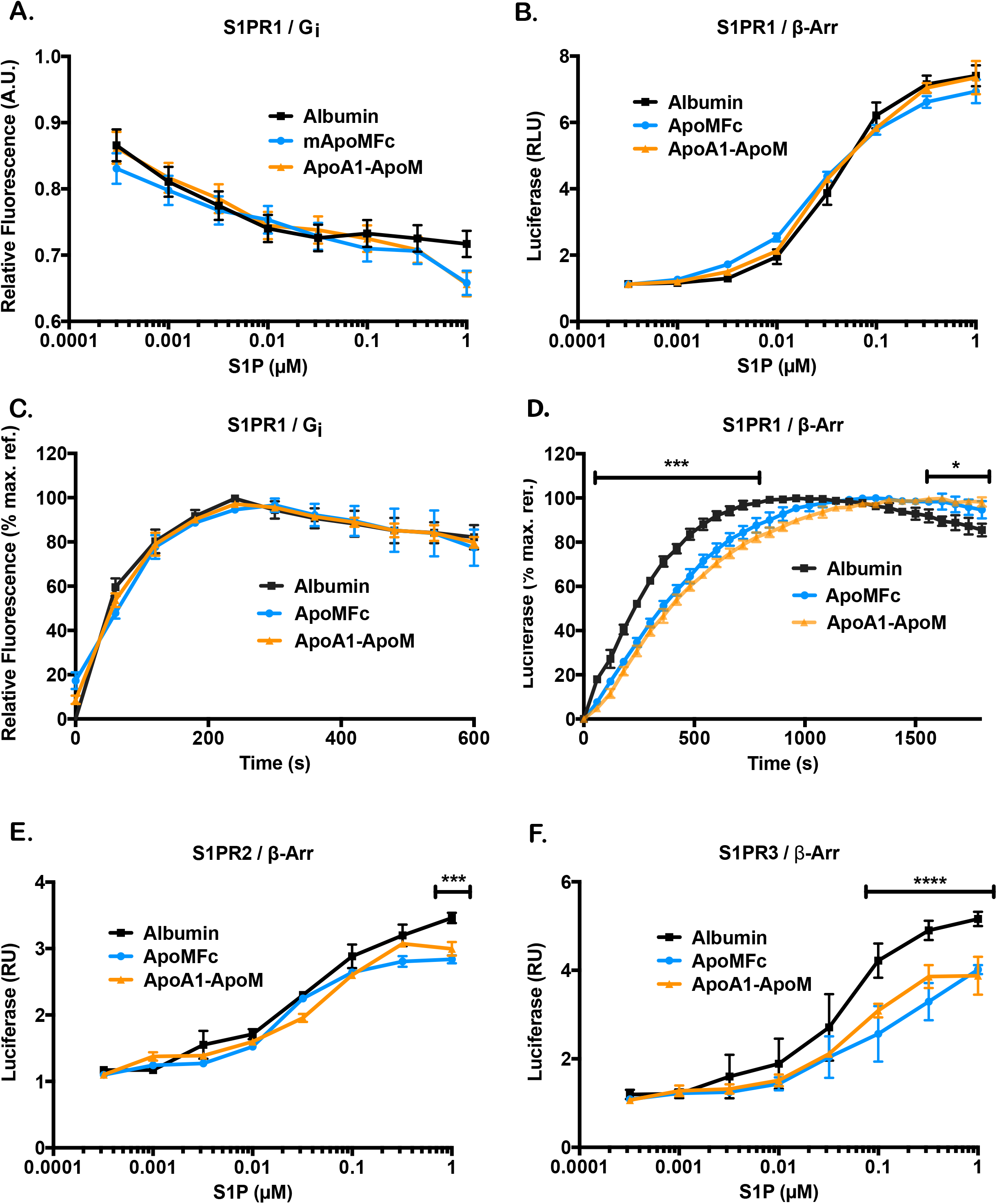
A1M/S1P activates S1PRs. NanoBiT analysis was performed to analyze of S1PR1 activation of Gα_i_ **(A)** and β-arrestin **(B)**. Albumin-S1P, ApoM-Fc/S1P, and A1M/S1P were assayed by dose-response analysis (0.32nM, 1nM, 3.2nM, 10nM, 32nM, 100nM, 320nM, 1 μM S1P). N = 3. Temporal analysis of S1P-dependent (1μM) receptor activation of G_i_α **(C)** and β-arrestin **(D).** N=3. Maximal peaks were analyzed by multiple paired student t-test (Albumin, P< 0.003; ApoM, P<0.03) NanoBiT analysis was performed to analyze S1PR2 and S1PR3 receptor coupling to β-arrestin **(E, F)** using the same dose-response as above. N = 3. Multiple paired student t-test analysis S1PR2 (1μM, P<0.005) and S1P3 (100nM-1μM, P<0.001).

We also examined the ability of the various chaperones to activate S1PR2 and S1PR3 (Figure 2E–F). We observed greater maximum stimulation of these receptors with albumin-S1P, when compared to either ApoM-Fc/S1P or A1M/S1P (P<0.005), suggesting that ApoM-bound S1P activates S1PR1 avidly in a Gα_i_-biased manner.

### A1M/S1P stimulates endothelial barrier function

To determine the ability of A1M/S1P regulate endothelial cell barrier function, trans-endothelial electrical resistance (TEER) analysis was performed on HUVEC after A1M/S1P stimulation (26). We performed a dose response study based on S1P content (S1P 10-300nM; A1M 0.8-24μg/ml) and observed a robust enhancement of endothelial barrier function (Figure 3A). Unloaded A1M did not have activity. Dose-response studies show equivalence of A1M/S1P and ApoM-Fc/S1P (EC_50_ ~ 2.5 μg/ml; 30nM S1P) suggesting the functionality of ApoM as a S1P chaperone whether it is in a soluble or nanodisc form (Figure 3B). As expected, unloaded A1M and ApoM-Fc did not stimulate the endothelial barrier function.

**Fig. 3.**
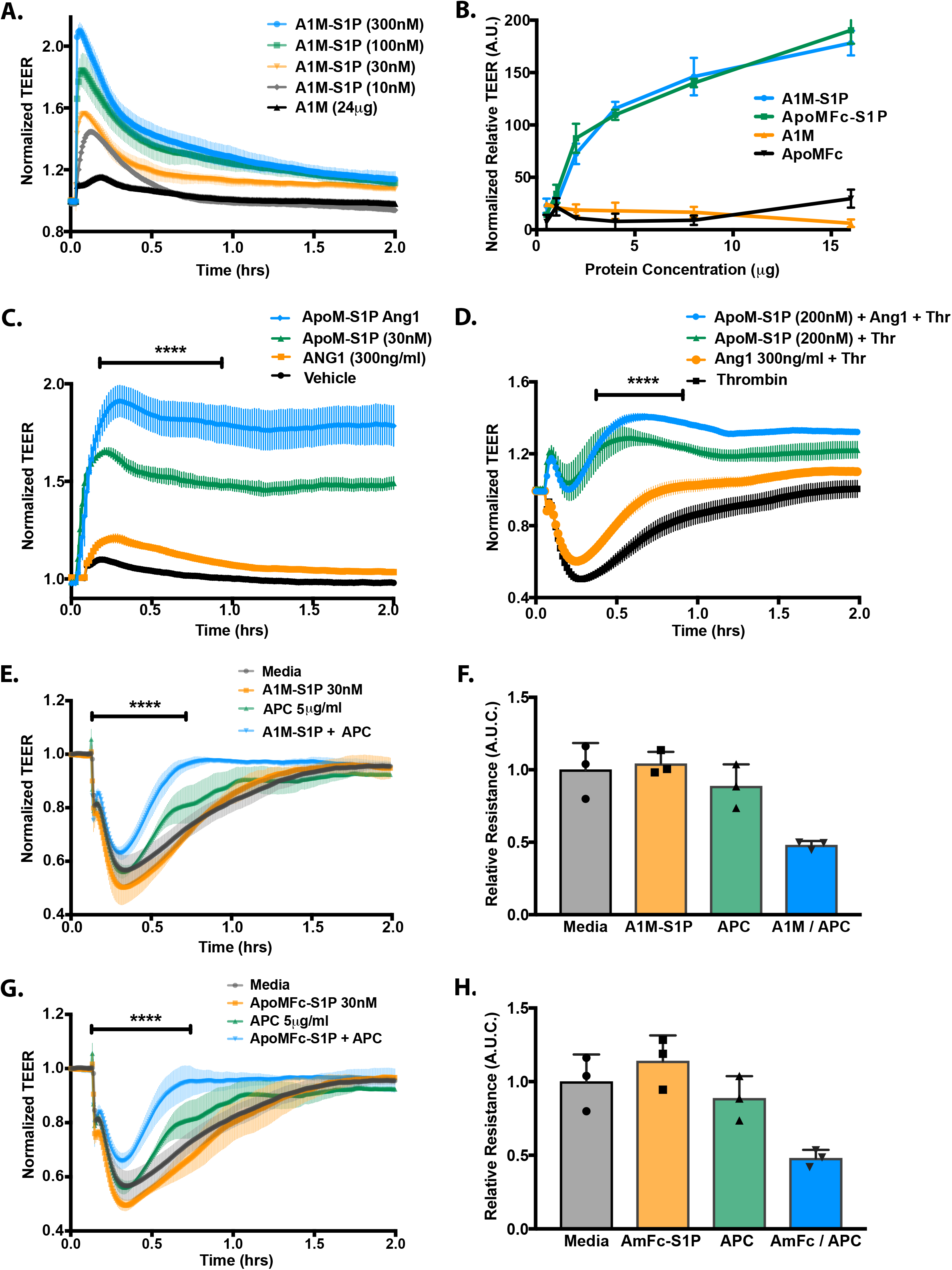
ApoA1-ApoM/S1P maintains endothelial barrier function *in vitro* and co-protects endothelial barrier in response to thrombin-induced barrier degradation. **(A)** A1M/S1P-dependent enhancement of barrier function in HUVEC (24 μg/ml A1M contains ~ 300 nM S1P). **(B)** Comparison of A1M/S1P and ApoM-Fc/S1P by TEER using ApoA1-ApoM/S1P (16μg/ml of A1M or ApoM-Fc ~ 200nM S1P). Unloaded chaperones were used as controls. Data are presented as area under the curve (N = 3; mean ± SD. P < 0.0001, two-way ANOVA followed by paired student t-test t test). **(C)** HUVECs were analyzed for barrier protection by TEER analysis using ApoM-Fc/S1P (30 nM) and Ang-1 (300 ng/ml) individually or in combination. **(D)** HUVECs were analyzed for barrier protection by TEER analysis in response to thrombin (1U/ml) treatment using ApoM-Fc/S1P (200nM), Ang-1 (300ng/ml) or in combination. For C and D, data were analyzed by non-parametric t-test (Mann-Whitney) (****P<0.0001). **(E-H)** HUVECs were analyzed for barrier protection by TEER analysis using either A1M/S1P (30nM) (E) or ApoM-Fc/S1P (30nM) (G) in conjunction with APC (5μg/ml). After 1 hour pretreatment, thrombin was added for an additional 2 hours. Area Under the Curve was analyzed (n=3) followed by non-parametric t-test (Mann-Whitney) of control or individual treatments vs combined treatments (F,G). ****P<0.0001 for (E,G) and P<0.01 for (F,G).

Angiopoietin-1 (Ang-1) is a polypeptide that stimulates endothelial barrier function and attenuates vascular leak during inflammation and thrombosis (37, 38). ApoM-Fc/S1P and Ang-1 enhanced endothelial barrier in an additive manner (Figure 3C). Thrombin, a protease produced during blood clotting (hemostasis), activates its GPCRs to disrupt endothelial barrier function. S1P is known to dampen thrombin-induced barrier breach by activating S1PRs (39). We performed a dose response study to evaluate the effect of ApoM-Fc/S1P on thrombin-induced barrier degradation and confirmed previous observations of S1P-dependent barrier protection against thrombin and found that S1P enhanced barrier function in a dose-dependent manner (Supplementary Figure 2A). ApoM-Fc/S1P and Ang-1 cooperated to suppress thrombin-induced barrier breach (Figure 3D). These data are consistent with S1P and Ang-1 protecting the endothelial barrier in homeostatic and inflammatory conditions cooperatively.

Activated Protein C (APC) antagonizes thrombin by interacting with endothelial cell surface receptors such as thrombomodulin and the endothelial protein C receptor (40). We evaluated APC in a thrombin-induced HUVEC barrier breach assay (Supplementary Figure 2B) and then in conjunction with both A1M/S1P (Figure 3E) and ApoM-Fc/S1P (Figure 3G), in all cases using sub-optimal concentrations of both APC (5μg/ml) and S1P (30nM). Area under the curve analysis revealed a 51-56% reversal of thrombin-induced barrier breach for A1M/S1P and a 47-58% rescue for ApoM-Fc/S1P (Figure 3F,H). Our results reveal that S1P bound to A1M or ApoM-Fc cooperates with APC to blunt thrombin-induced vascular leak, suggesting this combination may have utility in blocking the inflammatory effects of thrombin. Together these data reveal the cooperative interactions between endothelial barrier protective agents S1P, Ang-1 and APC.

### A1M binds to the stable prostacyclin analog Iloprost, which cooperates with S1P to enhance barrier function

Endothelial cell derived PGI_2_ acts the IP receptor to activate the Gα_s_/ adenylate cyclase/ cAMP pathway to suppress platelet aggregation, vascular smooth muscle dilitation and inflammatory responses (41). Recently, IP receptor activators were shown to suppress inflammation-induced vascular leak, likely due to the engagement of the cAMP/ EPAC/ Rap1 pathway. The lability of PGI_2_ due to autohydrolysis is inhibited by HDL association (17). Indeed, we found that purified A1M nanodiscs can associate with Iloprost, a stable PGI_2_ analog (42) (Figure 4A). Iloprost-bound to ApoA1 was able to activate the IP receptor, as determined by IP/ cAMP-responsive CREB luciferase reporter assay (Figure 4B).

**Fig. 4.**
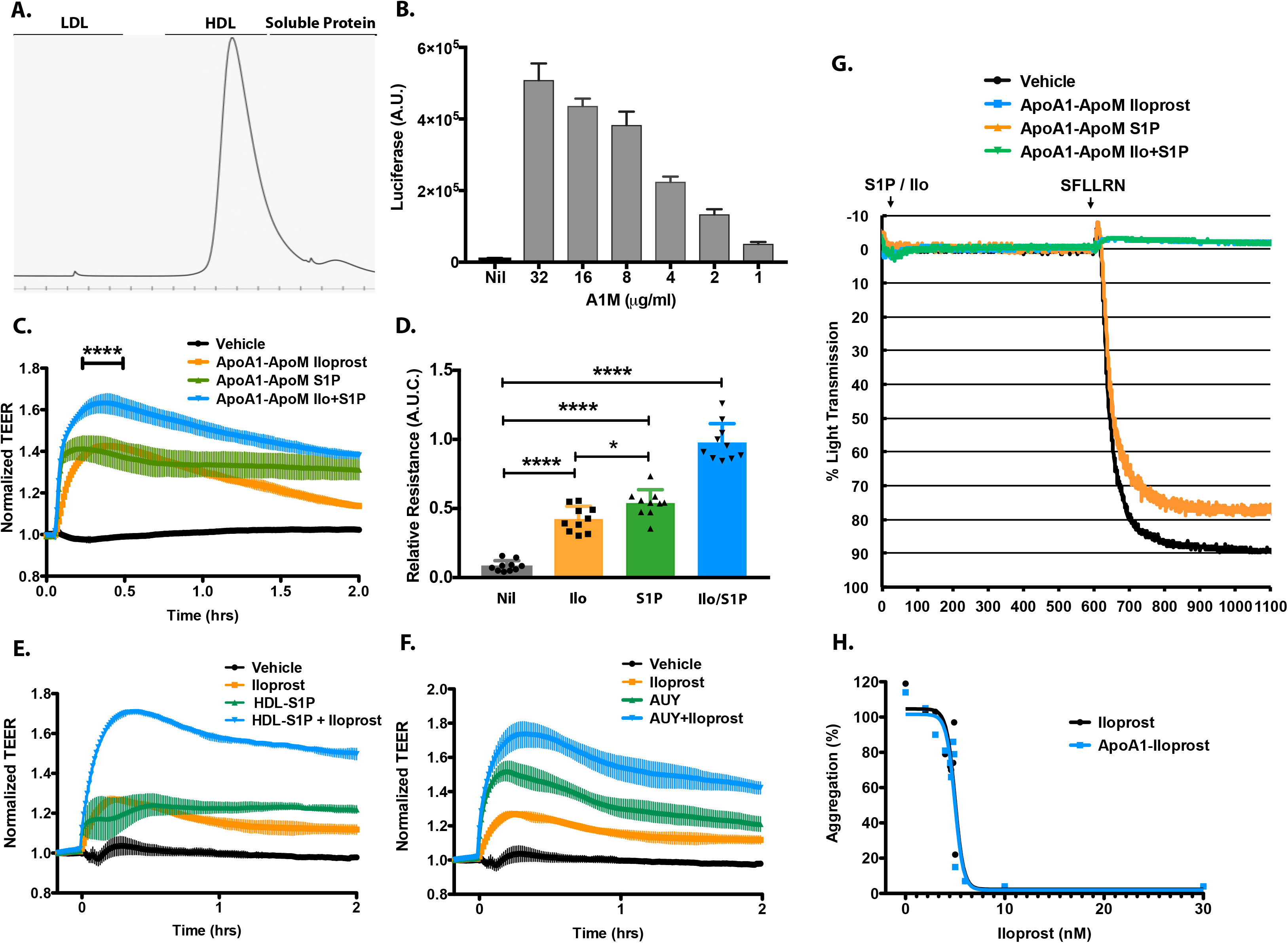
Production and characterization of A1M/Iloprost. Spectrophotometric (OD 280 nm) FPLC trace of A1M lipidated with PC and Iloprost (1 mg). CREB-luciferase reporter cell line was stimulated with vehicle or A1M/Iloprost for 8 hours and cell lysates were assayed for Luciferase activity. **(C,D)** A1M/S1P (8μg/ml; 100nM S1P), or A1M/Iloprost (25μg/ml; 200nM Iloprost) stimulate HUVEC barrier function. (N = 4) mean ± SEM. ****P < 0.0001, *P=0.05 based on two-way ANOVA followed by non-parametric t-test (Mann-Whitney). TEER analysis of HDL-S1P (100nM S1P) **(E)** and the S1PR1 agonist AUY954 (1μM) **(F)** alone, and in combination with Iloprost (200nM) (N ≥3). **(G)** Isolated human platelets were assayed for aggregation in response to the thrombin mimetic peptide SFLLRN. Presented are the results for replicate analyses for vehicle control, A1M/Iloprost (10 nM), A1M/S1P (200 nM), or in combination **(H).** A narrow-range dose response was performed using either free Iloprost or equivalent ApoA1/Iloprost (0, 2, 3, 4, 5, 6, 10, 30nM drug). N=2.

Since the cAMP/ EPAC/ Rap1 pathway stimulates endothelial barrier function, we tested A1M-bound Iloprost. As shown in Figure 4C, A1M/Iloprost stimulated endothelial barrier function to a similar degree as low dose A1M/S1P (S1P 30nM). Combination of A1M/Iloprost and A1M/S1P provided additive barrier promotion of endothelial cells (Figure 4C). Indeed unbound Iloprost (200nM) cooperated with HDL-S1P (Figure 4E) and the S1PR1-selective agonist AUY954 (Figure 4F) to enhance endothelial barrier function in an additive manner. These data are consistent with A1M stabilizing PGI_2_ such that it may protect the barrier function of the vascular endothelium in a cooperative manner with S1P.

Prostacyclin signaling through the IP receptor inhibits platelet aggregation thus achieving inhibition of thrombosis (43). A1M/Iloprosst potently inhibited platelet aggregation induced by the thrombin receptor activating peptide (Figure 4G). Dose-response studies indicated an EC_50_ of ~ 11 nM for free Iloprost and ~9 nM for ApoA1 nanodisc-associated-Iloprost (Figure 4H), suggesting a complete retention of drug potency when Iloprost is associated with HDL-like nanodiscs. In contrast, A1M/S1P did not inhibit platelet aggregation.

### ApoA1 attenuates TNFα-induced NF-κB activity and ICAM-1 expression

HDL-bound S1P is known to attenuate TNFα-induced ICAM-1 expression (21, 44). In contrast, albumin-bound S1P did not suppress ICAM-1 expression, suggesting that S1PR1 agonism alone is not sufficient to suppress cytokine inflammatory responses. Other studies have suggested that the ApoA1 moiety of HDL can engage endothelial cells to induce NO release (45) and suppress TLR4- and TNFα-induced NFκB activation in myeloid cells (14). Using a luciferase-based NF-κB transcriptional reporter assay, we evaluated whether ApoA1 or A1M + S1P treatment regulates cytokine-induced NFκB transcription and whether S1PR1 signaling could influence this action. As shown in Figure 5A, low doses of ApoA1, A1M and A1M/S1P (25-50 μg/ml) inhibited TNFα-induced NF-κB activity ~25%. Higher doses (100-400 μg/ml) were not as inhibitory and the dose-response curve exhibited a bi-phasic pattern. In contrast, ApoM-Fc/S1P and albumin-S1P did not inhibit TNFα-induced NF-κB activity (Figure 5B). Treatment of HUVEC with the cytokine TNF-α induced ICAM-1, A1M lipoprotein particles suppressed TNF-α induced ICAM-1 regardless of the bound lipid mediator (either S1P or Iloprost). None of the ApoM-Fc preparations were effective (Figure 5C). These data are consistent with antagonism of TNFα-induced endothelial inflammatory pathways by the ApoA1 moiety of the A1M particle.

**Fig. 5.**
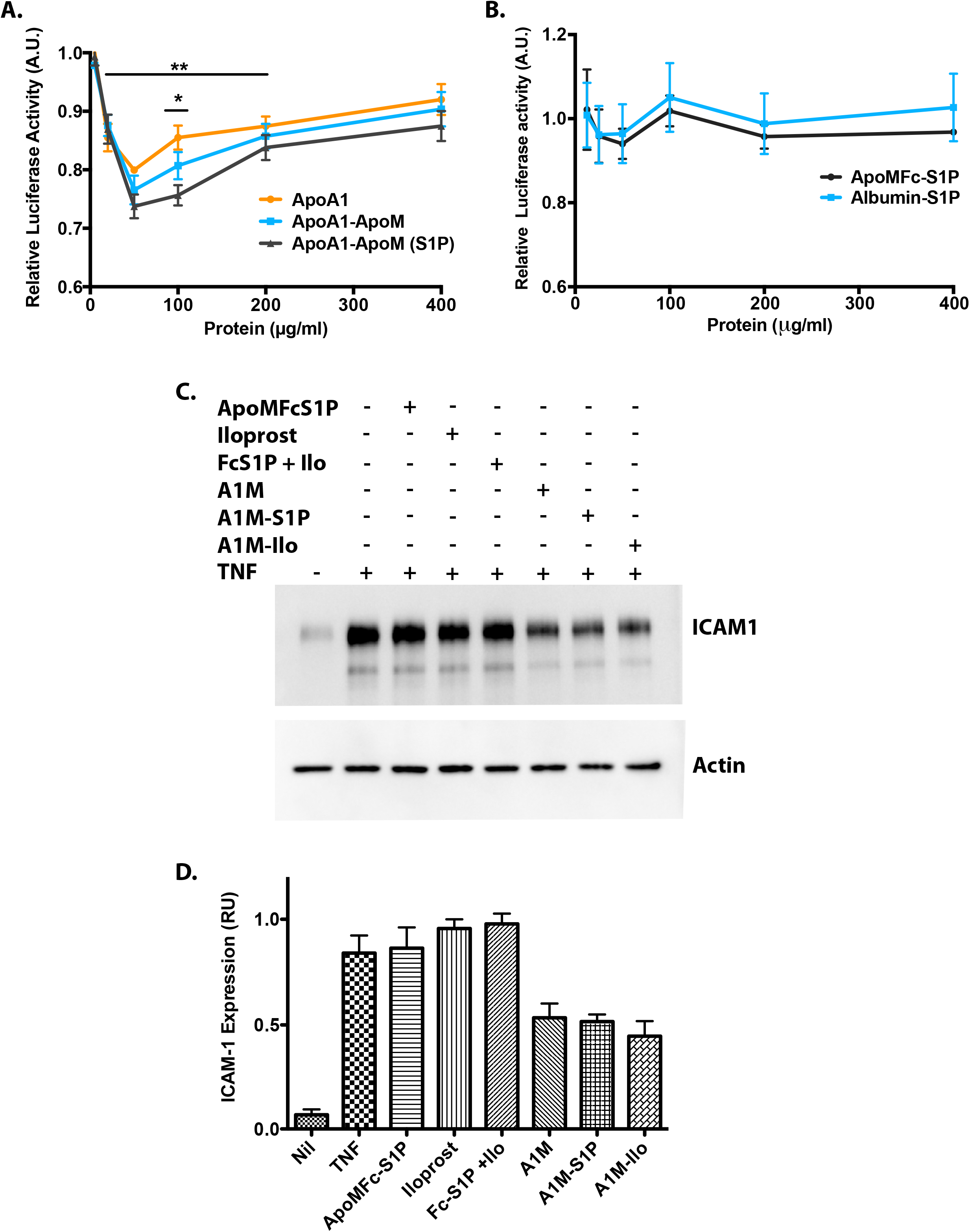
A1M attenuates TNFα-dependent Inflammation. **(A)** HMEC NF-κB-luciferase reporter cells were assayed for TNFα-induced NF-κB luciferase reporter activity in the presence of ApoA1, A1M and A1M/S1P. (N=3; Mean + S.D.; **P< 0.01 *P<0.05 Student t-test). **(B)** Effect of ApoM-Fc and Albumin-S1P on TNFα-induced NF-κB luciferase reporter activity (N= 3 Mean + S.D.). **(C)** HUVECs were assayed for TNFα induction of ICAM-1 expression by immunoblot analysis. Cultures were pre-treated for 10 minutes with ApomFc/S1P (100nM), Iloprost (200nM), or both in combination, as well as A1M, A1M/S1P, A1M/Iloprosst (all 200μg/ml) and induced with TNFα (10ng/ml) for 5 hours. Image J was used to obtain semi-quantitative values from scans of immunoblots (N=3).

### A1M/S1P suppresses inflammation

We next examined the anti-inflammatory actions of A1M/S1P on neutrophils, the most abundant cells of the innate immune system. Neutrophil activation results in an oxidative burst, which produces reactive oxygen species (ROS) such as superoxide anion (O_2_^−^), hydroxyl radical and singlet oxygen, which are damaging to cells and tissues (46). We isolated thioglycolate-elicted mouse neutrophils and tested their oxidative burst in response to formyl peptide (fMLF) activation. As shown in Figure 6A, neutrophil ROS generation was inhibited by both ApoM-S1P (~42%) or ApoA1/Iloprost (~44%) co-treatment. Combination treatment did not further decrease ROS formation. The ability of A1M-bound S1P and PGI_2_ to suppress neutrophil oxidant stress is consistent with HDL attenuating thrombo-inflammatory reactions via its bioactive lipid mediator cargo (S1P and PGI_2_).

**Fig. 6.**
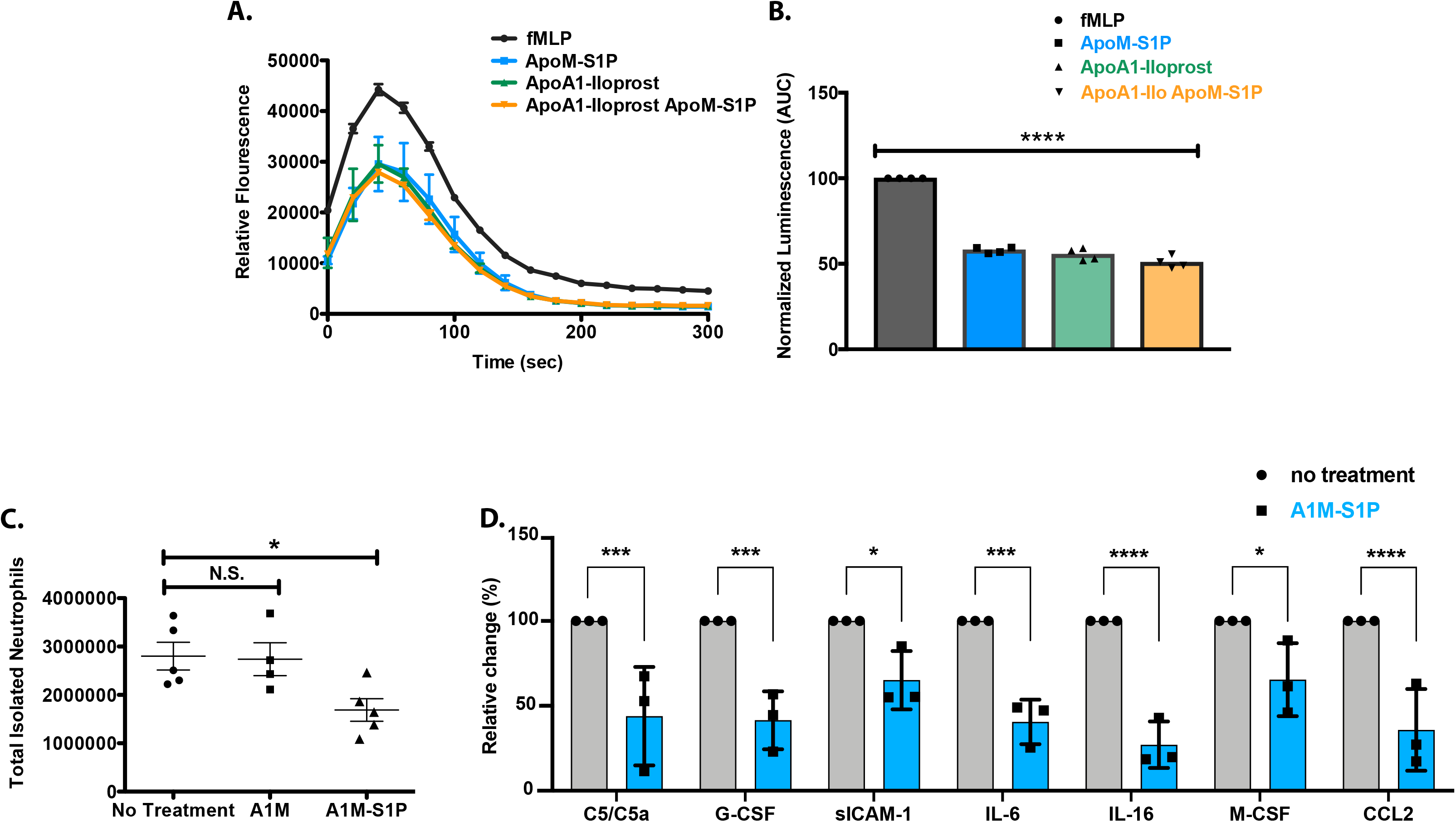
Suppression of neutrophil activation by A1M and related chaperones. **(A)** Isolated peritoneal Mouse Neutrophils were assayed for oxidative burst after fMLF stimulation. Cells were pre-incubated for 10 minutes with vehicle control, ApoM-Fc/S1P (S1P 100nM), ApoA1/Iloprost (160nM), or in combination. Data are presented as normalized luminol fluorescence **(B)** Composite data for N=4 independent experiments. Area under the Curve (AUC) data were analyzed by non-parametric t-test (Mann-Whitney) and P values <0.0001. **(C)** Mice were treated with with thioglycolate and either PBS, A1M (200μg) or A1M/S1P (200μg). Peritoneal cells (4 h) were counted and analyzed by flow cytometry. Data were analyzed by one-way ANOVA followed by student t-test P=0.0317. **(D)** Peritoneal lavage supernatant from (C) was assayed by cytokine array analysis. Resulting blots were analyzed on IMAGEJ and differentially expressed analytes were quantified and data are expressed after normalization (Control = 100%). Data was analyzed by 2-way ANOVA followed by multiple paired student t-test (C5/C5a P= 0.028, G-CSF P= 0.004, sICAM-1 P= 0.025, IL-6 P= 0.001, IL-16 P= 0.0007, M-CSF P= 0.049, CCL2 P=0.009).

We next examined if A1M/S1P can impact inflammatory responses in a murine sterile inflammation model. Thioglycolate injection into the peritoneum induces rapid neutrophil-influx and inflammation. We found that A1M/S1P injected one hour after the initiation of the inflammatory response markedly suppressed neutrophil influx into the peritoneum at 4 h. S1P association is needed for A1M to suppress neutrophil infiltration. Furthermore, peritoneal lavage fluid contained reduced immunoreactivity of inflammatory factors - complement C5a, G-CSF, soluble ICAM-1, IL-6, IL-16, M-CSF and CCL2 in A1M/S1P treated animals. These data provide evidence for an anti-inflammatory function of A1M/S1P in vivo.

## Discussion

The induction of vascular dysfunction during infectious, traumatic, or chronic metabolic illnesses, and aging has been identified as a primary event for the rise of subsequent tissue and organ pathology. Disrupted and inflamed vasculature are edematous and trigger both platelet-driven thrombosis and the subsequent activation of the innate immune response, principally through neutrophils, cumulatively leading to greater organ degradation. Extensive studies of HDL have revealed multiple vascular protective properties that ameliorate these pathological processes (9, 10, 16). Both nascent and lipidated ApoA1, the principal structural protein of HDL, induce reverse cholesterol efflux, which can reduce inflammatory cytokine driven signal transduction (47). In addition, HDL was found to induce and chaperone the anti-inflammatory lipid prostacyclin, which acts to inhibit platelet activation and thrombosis, block neutrophil adherence to endothelium, and enhance endothelial barrier function (48). Finally, recent work identified the HDL associated protein, Apolipoprotein M, as the principal chaperone of the biologically active sphingolipid, S1P, which enhances endothelial homeostasis and barrier function (19, 49).

To take advantage of these vascular protective aspects of HDL, we created a recombinant fusion protein, ApoA1-ApoM (A1M), which incorporates the identified anti-inflammatory properties of ApoA1 as well as the barrier protecting properties of S1P. A1M/S1P was demonstrated to activate both Giα and β-arrestin coupling in the three endothelial S1P receptors, S1PR1, S1PR2, and S1PR3 and regulate endothelial barrier function as judged by TEER analysis, as we have reported for other S1P chaperones (26). Both A1M and A1M/S1P were anti-inflammatory, reducing both TNFα-induced NF-κB activation in a reporter assay and inhibiting downstream inflammatory ICAM1 expression in endothelium. Notably, we observed that other S1P chaperones ApoM-Fc and albumin had no inhibitory activity, suggesting that the ApoA1 moiety is the primary inhibitor of TNFR signaling.

We lipidated the ApoA1 moiety in vitro to enhance the potential function of A1M, using standard procedures and demonstrated that the stable PGI_2_ analogue, Iloprost, can be functionally incorporated into A1M, creating A1M/Iloprosst. We demonstrated that A1M/Iloprosst retained the established properties of PGI_2_ including activation of PKA-dependent CREB-signaling, and enhancement of endothelial barrier protection. Intriguingly, in further studies, we demonstrated that both free and A1M-bound Iloprost could provide additive protection of endothelial barrier function in conjunction with A1M/S1P, suggesting a useful co-operation of these two pathways for endothelial function and protection.

Initial activation of the endothelium activates multiple pathways that result in further amplification of the inflammatory process including thrombin formation, which induces disruption of the endothelial barrier and subsequent platelet activation leading to thrombosis (16). We demonstrated that A1M/S1P blocks thrombin-induced barrier disruption and enhances the barrier-protective activity of APC. Moreover, we showed that A1M/Iloprosst inhibits thrombin-driven platelet aggregation in vitro. A second consequence of endothelial inflammation is neutrophil recruitment and activation leading to the generation of ROS, a destructive mechanism induced by bacterial infections, tissue damage and inflammatory cytokines (2). In this regard, we demonstrated the potential utility of A1M/Iloprosst/S1P, observing that both lipid ligands potently inhibited the generation of ROS in isolated neutrophils. Our approach shows the potential of dampening both platelet-driven thrombosis and neutrophil-dependent tissue damage during tissue inflammation.

These results taken together argue that our recombinant A1M/S1P/Iloprost has the potential to provide a three-pronged approach to recapitulate the established protective effects of HDL by enhancing endothelial function with S1P, inhibiting inflammatory cytokine signaling with ApoA1, and inhibiting the amplification of inflammation by dampening platelet and neutrophil activation via PGI_2_. This approach has potential utility in many inflammatory states in which endothelial damage driven by infectious pathogens, vascular reperfusion injury after trauma or in organ transplantation, as well as more chronic vasculopathies associated with atherosclerosis, diabetes and autoimmunity.

## Supplementary Figure Legends

**Supplementary Figure 1. Nucleotide and Amino acid sequence of ApoA1-ApoM identifying the features of the fusion protein.**

The ApoA1-ApoM fusion contains the signal peptide (highlighted in yellow) of native murine ApoA1 and amino acids 1-264 of the open reading frame followed by one (Gly4Ser1) flexible linker region, amino acids 21-190 of murine ApoM, a 6x histidine purification tag, and a stop translation codon.

**Supplementary Figure 2 Titration assays for Thrombin inhibition by TEER analysis**

**(A)** HUVECs were analyzed for barrier protection by TEER analysis (n=3) for 2 hours using ApoM-S1P (1, 3, 10, 30, and 100nM S1P) in combination with Thrombin (1U/ML). (B) HUVECs were analyzed for barrier protection by TEER analysis (n=3) for 2 hours using Activated Protein C (APC; 1, 5, 20, and 50μg/ml) in combination with Thrombin (1U/ml).

## Acknowledgements

This work was funded by NIH grants R35HL135821 and R01EY031715 to T.H. A.I. was funded by Japan Society for the Promotion of Science (JSPS) KAKENHI grants 21H04791, 21H05113 and JPJSBP120218801; Moonshot Research and Development Program JPMJMS2023 and FOREST Program JPMJFR215T from Japan Science and Technology Agency (JST); the LEAP JP20gm0010004 and the BINDS JP20am0101095 from the Japan Agency for Medical Research and Development (AMED); Ono Medical Research Foundation; Takeda Science Foundation.”

## REFERENCES

1. J. S. Pober, W. Min, Endothelial cell dysfunction, injury and death. Handb Exp Pharmacol 10.1007/3-540-36028-x_5, 135–156 (2006).

2. J. S. Pober, W. C. Sessa, Evolving functions of endothelial cells in inflammation. Nat Rev Immunol 7, 803–815 (2007).

3. P. Castro et al., Is the Endothelium the Missing Link in the Pathophysiology and Treatment of COVID-19 Complications? Cardiovasc Drugs Ther 10.1007/s10557-021-07207-w (2021).

4. R. Y. Kiseleva et al., Targeting therapeutics to endothelium: are we there yet? Drug Deliv Transl Res 8, 883–902 (2018).

5. L. Yvan-Charvet, N. Wang, A. R. Tall, Role of HDL, ABCA1, and ABCG1 transporters in cholesterol efflux and immune responses. Arterioscler Thromb Vasc Biol 30, 139–143 (2010).

6. M. Ouimet, T. J. Barrett, E. A. Fisher, HDL and Reverse Cholesterol Transport. Circ Res 124, 1505–1518 (2019).

7. A. Ossoli, A. T. Remaley, B. Vaisman, L. Calabresi, M. Gomaraschi, Plasma-derived and synthetic high-density lipoprotein inhibit tissue factor in endothelial cells and monocytes. Biochem J 473, 211–219 (2016).

8. A. T. Remaley, G. D. Norata, A. L. Catapano, Novel concepts in HDL pharmacology. Cardiovasc Res 103, 423–428 (2014).

9. A. Rohatgi, M. Westerterp, A. von Eckardstein, A. Remaley, K. A. Rye, HDL in the 21st Century: A Multifunctional Roadmap for Future HDL Research. Circulation 143, 2293–2309 (2021).

10. A. C. Sposito et al., HDL-Targeted Therapies During Myocardial Infarction. Cardiovasc Drugs Ther 33, 371–381 (2019).

11. J. W. Heinecke, The protein cargo of HDL: implications for vascular wall biology and therapeutics. J Clin Lipidol 4, 371–375 (2010).

12. B. Shao, J. W. Heinecke, Quantifying HDL proteins by mass spectrometry: how many proteins are there and what are their functions? Expert Rev Proteomics 15, 31–40 (2018).

13. M. Suzuki et al., High-density lipoprotein suppresses the type I interferon response, a family of potent antiviral immunoregulators, in macrophages challenged with lipopolysaccharide. Circulation 122, 1919–1927 (2010).

14. P. Fotakis et al., Anti-Inflammatory Effects of HDL (High-Density Lipoprotein) in Macrophages Predominate Over Proinflammatory Effects in Atherosclerotic Plaques. Arterioscler Thromb Vasc Biol 39, e253–e272 (2019).

15. C. Mineo, H. Deguchi, J. H. Griffin, P. W. Shaul, Endothelial and antithrombotic actions of HDL. Circ Res 98, 1352–1364 (2006).

16. C. Mineo, P. W. Shaul, Regulation of signal transduction by HDL. J Lipid Res 54, 2315–2324 (2013).

17. H. Morishita, Y. Yui, R. Hattori, T. Aoyama, C. Kawai, Increased hydrolysis of cholesteryl ester with prostacyclin is potentiated by high density lipoprotein through the prostacyclin stabilization. J Clin Invest 86, 1885–1891 (1990).

18. Y. Yui et al., Serum prostacyclin stabilizing factor is identical to apolipoprotein A-I (Apo A-I). A novel function of Apo A-I. J Clin Invest 82, 803–807 (1988).

19. C. Christoffersen et al., Endothelium-protective sphingosine-1-phosphate provided by HDL-associated apolipoprotein M. Proc Natl Acad Sci U S A 108, 9613–9618 (2011).

20. R. L. Proia, T. Hla, Emerging biology of sphingosine-1-phosphate: its role in pathogenesis and therapy. J Clin Invest 125, 1379–1387 (2015).

21. S. Galvani et al., HDL-bound sphingosine 1-phosphate acts as a biased agonist for the endothelial cell receptor S1P1 to limit vascular inflammation. Sci Signal 8, ra79 (2015).

22. V. A. Blaho et al., HDL-bound sphingosine-1-phosphate restrains lymphopoiesis and neuroinflammation. Nature 523, 342–346 (2015).

23. Z. Cao, Guo, P., Yang, D., Swendeman, S.L., Wang, Z., Christoffersen, C., Nielsen, L.B., Friedman, S.L., Powell, C.A., Hla, T., Ding, B., Aging attenuates interorgan protective signaling axis mediated by ApoM-bound sphingosine 1-phosphate. Dev Cell In Press (2020).

24. B. S. Ding et al., HDL activation of endothelial sphingosine-1-phosphate receptor-1 (S1P1) promotes regeneration and suppresses fibrosis in the liver. JCI Insight 1, e87058 (2016).

25. S. Velagapudi et al., Apolipoprotein M and Sphingosine-1-Phosphate Receptor 1 Promote the Transendothelial Transport of High-Density Lipoprotein. Arterioscler Thromb Vasc Biol 41, e468–e479 (2021).

26. S. L. Swendeman et al., An engineered S1P chaperone attenuates hypertension and ischemic injury. Sci Signal 10 (2017).

27. N. Burg, S. Swendeman, S. Worgall, T. Hla, J. E. Salmon, Sphingosine 1-Phosphate Receptor 1 Signaling Maintains Endothelial Cell Barrier Function and Protects Against Immune Complex-Induced Vascular Injury. Arthritis Rheumatol 70, 1879–1889 (2018).

28. M. J. Lee et al., Sphingosine-1-phosphate as a ligand for the G protein-coupled receptor EDG-1. Science 279, 1552–1555 (1998).

29. H. Ru et al., Molecular Mechanism of V(D)J Recombination from Synaptic RAG1-RAG2 Complex Structures. Cell 163, 1138–1152 (2015).

30. E. Engelbrecht et al., Sphingosine 1-phosphate-regulated transcriptomes in heterogenous arterial and lymphatic endothelium of the aorta. Elife 9 (2020).

31. A. Schwendeman et al., The effect of phospholipid composition of reconstituted HDL on its cholesterol efflux and anti-inflammatory properties. J Lipid Res 56, 1727–1737 (2015).

32. M. N. Oda et al., Reconstituted high density lipoprotein enriched with the polyene antibiotic amphotericin B. J Lipid Res 47, 260–267 (2006).

33. T. Hla, K. Neilson, Human cyclooxygenase-2 cDNA. Proc Natl Acad Sci U S A 89, 7384–7388 (1992).

34. Y. Hisano et al., Lysolipid receptor cross-talk regulates lymphatic endothelial junctions in lymph nodes. J Exp Med 216, 1582–1598 (2019).

35. J. Michaud, M. Kohno, R. L. Proia, T. Hla, Normal acute and chronic inflammatory responses in sphingosine kinase 1 knockout mice. FEBS Lett 580, 4607–4612 (2006).

36. R. Trinh, B. Gurbaxani, S. L. Morrison, M. Seyfzadeh, Optimization of codon pair use within the (GGGGS)3 linker sequence results in enhanced protein expression. Mol Immunol 40, 717–722 (2004).

37. M. Jeansson et al., Angiopoietin-1 is essential in mouse vasculature during development and in response to injury. J Clin Invest 121, 2278–2289 (2011).

38. S. M. Parikh, Angiopoietins and Tie2 in vascular inflammation. Curr Opin Hematol 24, 432–438 (2017).

39. J. G. Garcia et al., Sphingosine 1-phosphate promotes endothelial cell barrier integrity by Edg-dependent cytoskeletal rearrangement. J Clin Invest 108, 689–701 (2001).

40. J. H. Finigan et al., Activated protein C mediates novel lung endothelial barrier enhancement: role of sphingosine 1-phosphate receptor transactivation. J Biol Chem 280, 17286–17293 (2005).

41. H. Pluchart, C. Khouri, S. Blaise, M. Roustit, J. L. Cracowski, Targeting the Prostacyclin Pathway: Beyond Pulmonary Arterial Hypertension. Trends Pharmacol Sci 38, 512–523 (2017).

42. W. Skuballa, H. Vorbruggen, Synthesis of ciloprost (ZK 36 374): a chemically stable and biologically potent prostacyclin analog. Adv Prostaglandin Thromboxane Leukot Res 11, 299–305 (1983).

43. S. Moncada, E. A. Higgs, J. R. Vane, Human arterial and venous tissues generate prostacyclin (prostaglandin x), a potent inhibitor of platelet aggregation. Lancet 1, 18–20 (1977).

44. G. W. Cockerill, K. A. Rye, J. R. Gamble, M. A. Vadas, P. J. Barter, High-density lipoproteins inhibit cytokine-induced expression of endothelial cell adhesion molecules. Arterioscler Thromb Vasc Biol 15, 1987–1994 (1995).

45. I. S. Yuhanna et al., High-density lipoprotein binding to scavenger receptor-BI activates endothelial nitric oxide synthase. Nat Med 7, 853–857 (2001).

46. S. Kumar, M. Dikshit, Metabolic Insight of Neutrophils in Health and Disease. Front Immunol 10, 2099 (2019).

47. J. Robert, E. Osto, A. von Eckardstein, The Endothelium Is Both a Target and a Barrier of HDL’s Protective Functions. Cells 10 (2021).

48. A. A. Birukova et al., Iloprost improves endothelial barrier function in lipopolysaccharide-induced lung injury. Eur Respir J 41, 165–176 (2013).

49. P. M. Christensen et al., Impaired endothelial barrier function in apolipoprotein M-deficient mice is dependent on sphingosine-1-phosphate receptor 1. FASEB J 30, 2351–2359 (2016).

