## SupplementalFigures for "Engineered high-density lipoprotein particles that chaperone bioactive lipid mediators to combat endothelial dysfunction and thromboinflammation"

#### Sup. Fig 1

**A.**

ATGAAAGCTGTGGTGCTGGCCGTGGCTCTGGTCTTCCTGACAGGGAGCCAGGCTTGGCACGTATGGCAGCA
AGATGAACCCCAGTCCCAATGGGACAAAGTGAAGGATTTCGCTAATGTGTATGTGGATGCGGTCAAAGACA
GCGGCAGAGACTATGTGTCCCAGTTTGAATCCTCCTCCTTGGGCCAACAGCTGAACCTGAATCTCCTGGAA
AACTGGGACACTCTGGGTTCAACCGTTAGTCAGCTGCAGGAACGGCTGGGCCCATTGACTCGGGACTTCTG
GGATAACCTGGAGAAAGAAACAGATTGGGTGAGACAGGAGATGAACAAGGACCTAGAGGAAGTGAAACA
GAAGGTGCAGCCCTACCTGGACGAATTCCAGAAGAAATGGAAAGAGGATGTGGAGCTCTACCGCCAGAAG
GTGGCGCCTCTGGGCGCCGAGCTGCAGGAGAGCGCGCGCCAGAAGCTGCAGGAGCTGCAAGGGAGACTG
TCCCCTGTGGCTGAGGAATTTCGCGACCGCATGCGCACACACGTAGACTCTCTGCGCACACAGCTAGCGCC
CCACAGCGAACAGATGCGCGAGAGCCTGGCCCAGCGCCTGGCTGAGCTCAAGAGCAACCCTACCTTGAAC
GAGTACCACACCAGGGCCAAAACCCACCTGAAGACACTTGGCGAGAAAGCCAGACCTGCGCTGGAGGACC
TGCGCCATAGTCTGATGCCCATGCTGGAGACGCTTAAGACCAAAGCCCAGAGTGTGATCGACAAGGCCAG
CGAGACTCTGACTGCCCAGACGCGTACGCGGCCGCTTGGTGGAGGTGGATCTAATCAGTGCCCTGAGCAC
AGTCAACTAACTGCGCTGGGAATGGACGACACAGAGACCCCAGAGCCCCACCTGGGCCTGTGGTACTTTAT
TGCGGGAGCAGCCTCCACCACGGAAGAGTTGGCAACTTTTGATCCGGTGGACAATATTGTCTTCAACATGG
CTGCCGGCTCTGCCCCAAGGCAGCTCCAGCTTCGTGCTACCATCCGCACGAAAAGTGGGGTCTGTGTGCCC
CGGAAGTGGACATACCGATTGACTGAAGGGAAAGGAAACATGGAACTCAGAACTGAAGGGCGCCCAGACA
TGAAAACAGACCTGTTCTCCAGCTCGTGCCCAGGAGGAATCATGCTGAAAGAGACGGGCCAGGGCTACCA
GCGCTTTCTCCTCTACAATCGGTCACCACACCCTCCAGAGAAGTGTGTGGAGGAATTCCAGTCTCTGACCTC
TTGCTTGGACTTCAAAGCCTTCTTAGTGACTCCCAGGAATCAAGAGGCCTGCCCGCTGTCCAGCAAGCATC
ACCATCACCATCACTGA

**B.**

M K A V V L A V A L V F L T G S Q A W H V W Q Q D E P Q S Q W D K V K D F A N V Y V
D A V K D S G R D Y V S Q F E S S S L G Q Q L N L N L L E N W D T L G S T V S Q L Q E
R L G P L T R D F W D N L E K E T D W V R Q E M N K D L E E V K Q K V Q P Y L D E F
Q K K W K E D V E L Y R Q K V A P L G A E L Q E S A R Q K L Q E L Q G R L S P V A E
E F R D R M R T H V D S L R T Q L A P H S E Q M R E S L A Q R L A E L K S N P T L N E
Y H T R A K T H L K T L G E K A R P A L E D L R H S L M P M L E T L K T K A Q S V I D
K A S E T L T A Q T R T R P L G G G G S N Q C P E H S Q L T A L G M D D T E T P E P H
L G L W Y F I A G A A S T T E E L A T F D P V D N I V F N M A A G S A P R Q L Q L R A T
I R T K S G V C V P R K W T Y R L T E G K G N M E L R T E G R P D M K T D L F S S S C
P G G I M L K E T G Q G Y Q R F L L Y N R S P H P P E K C V E E F Q S L T S C L D F K
A F L V T P R N Q E A C P L S S K H H H H H H Stop

### Sup. Fig 2. TEER analyses of ApoM-S1P or APC after Thrombin treatment

A.

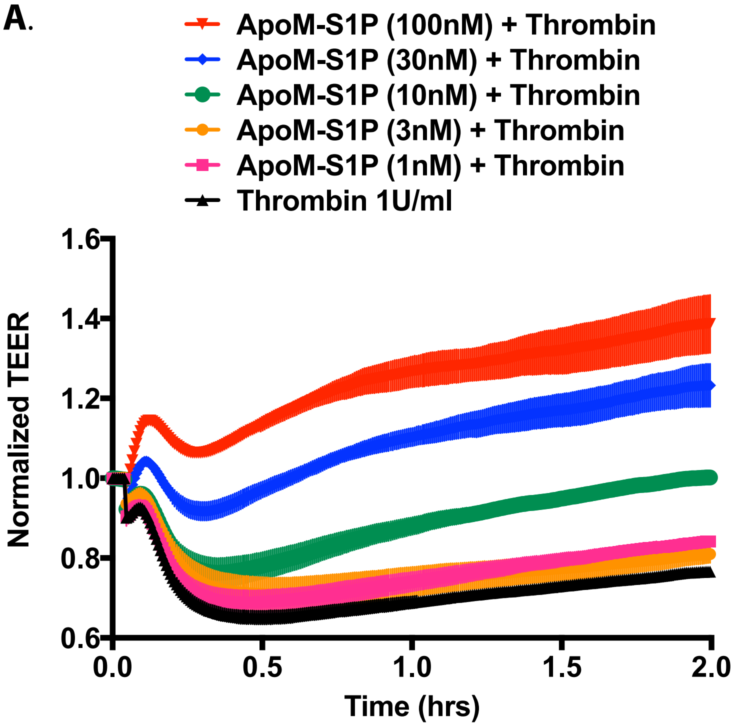


B.

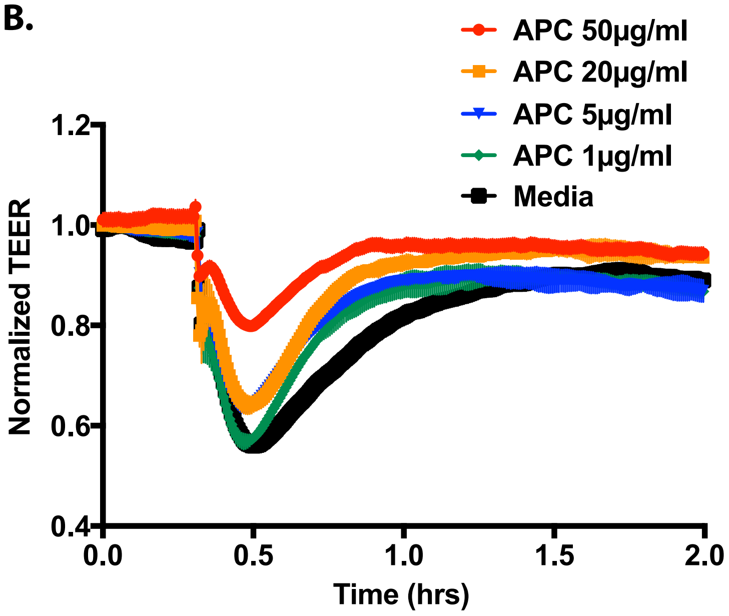
